# A cortico-cortical pathway targets inhibitory interneurons and modulates paw movement during locomotion in mice

**DOI:** 10.1101/2021.09.23.461507

**Authors:** Chia-wei Chang, Meiling Zhao, Samantha Grudzien, Max Oginsky, Yexin Yang, Sung Eun Kwon

**Affiliations:** Department of Molecular, Cellular, and Developmental Biology, University of Michigan, Ann Arbor, MI 48109; Department of Psychology, _UCLA_, Los Angeles, CA 90095; Neuroscience Graduate Program, University of Michigan, Ann Arbor, MI 48109

## Abstract

The primary somatosensory cortex (S1) is important for the control of movement as it encodes sensory input from the body periphery and external environment during ongoing movement. Mouse S1 consists of several distinct sensorimotor subnetworks that receive topographically organized cortico-cortical inputs from distant sensorimotor areas, including the secondary somatosensory cortex (S2) and primary motor cortex (M1). The role of the vibrissal S1 area and associated cortical connections during active sensing is well documented, but whether (and if so, how) non-whisker S1 areas are involved in movement control remains relatively unexplored. Here, we demonstrate that unilateral silencing of the non-whisker S1 area in both male and female mice disrupts hind paw movement during locomotion on a rotarod and a runway. S2 and M1 provide major long-range inputs to this S1 area. Silencing S2 → non-whisker S1 projections alters the hind paw orientation during locomotion while manipulation of the M1 projection has little effect. Using patch-clamp recordings in brain slices from male and female mice, we show that S2 projection preferentially innervates inhibitory interneuron subtypes. We conclude that interneuron-mediated S2–S1 cortico-cortical interactions are critical for efficient locomotion.

**Significance:** Somatosensory cortex participates in controlling rhythmic movements such as whisking and walking, but the neural circuitry underlying movement control by somatosensory cortex remains relatively unexplored. We uncover a cortico-cortical circuit in primary somatosensory cortex that regulates paw orientation during locomotion in mice. We identify neuronal elements that comprise these cortical pathways using pharmacology, behavioral assays and circuit-mapping methods.

## Introduction

The control of movement relies on a complex neural process involving multiple motor and sensory areas in the brain. The primary somatosensory cortex (S1) represents and processes self-generated (reafferent) and extrinsic (exafferent) inputs (Kaas et al., 1979, Chapin and Lin, 1984, O’Connor et al., 2013, Kim et al., 2015), thus playing a pivotal role in the control of movement and motor learning (Pavlides et al., 1993, Vidoni et al., 2010, Mathis et al., 2017, Karadimas et al., 2020). Abnormal processing of somatosensory information by S1 may lead to the motor deficits seen in neurological conditions typically classified as movement disorders (Tamura et al., 2009, Konczak et al., 2012). However, the neural circuitry underlying movement control by S1 remains relatively unexplored.

Mouse S1 consists of at least four distinct subregions that function as specialized somatosensory modules: the orofaciopharyngeal, upper limb, lower limb/trunk and whisker modules (Zingg et al., 2014). Each module forms a somatic sensorimotor subnetwork with sub- regions of the primary motor cortex (M1) and secondary somatosensory cortex (S2) via topographically arranged, reciprocal connections (Zingg et al., 2014). The whisker cortical sensorimotor subnetwork, including the S1 whisker area (wS1), has been extensively characterized in terms of anatomy, synaptic connectivity and function (Aronoff et al., 2010, Mao et al., 2011, Petreanu et al., 2012, Xu et al., 2012, Chen et al., 2013, Lee et al., 2013, Zagha et al., 2013, Clancy et al., 2015, Petrof et al., 2015, Zuo et al., 2015, Kinnischtzke et al., 2016, Kwon et al., 2016, Yamashita and Petersen, 2016, Zhang and Bruno, 2019, Naskar et al., 2021). Interactions between wS1 and other cortical areas, such as M1 and S2, are important for initiating whisking and active sensing during whisking (Petreanu et al., 2012, Xu et al., 2012, Lee et al., 2013, Zagha et al., 2013, Sreenivasan et al., 2016). These cortico-cortical interactions also underlie tactile perception (Chen et al., 2013, Zuo et al., 2015, Kwon et al., 2016, Yamashita and Petersen, 2016) and obstacle avoidance during locomotion (Warren et al., 2021).

Although coordinated movement must depend on inputs beyond whiskers, sensorimotor subnetworks beyond wS1 remain poorly characterized. Accumulating evidence suggests that S1 is involved in regulating locomotion. Changes in S1 neuronal activity correlate with locomotor output in primates (Fitzsimmons et al., 2009), cats (Favorov et al., 2015) and rodents (Chapin and Woodward, 1982a). More recently, a direct role for S1 in locomotion has been demonstrated (Karadimas et al., 2020). However, the specific aspects of locomotion that require functional S1 remain unclear. For example, cortical inputs to non-whisker S1 areas, including the limb/trunk subregion, may control rhythmic movement such as locomotion. The synaptic connectivity of different cortical inputs, and whether they serve distinct functions during locomotion, is unknown.

Here, we test the role(s) of mouse S1 and its long-range cortical inputs in locomotion and characterize their connectivity. We focus on the anteromedially located non-whisker subregion of S1 (nwS1), which corresponds to limb/trunk and dysgranular areas (Paxinos and Franklin, 2012). Using simple motor behavior assays and a pharmacological approach, we demonstrate that nwS1 activity is critical for maintaining hind paw orientation during locomotion. We identify S2 and M1 as major sources of cortical input to nwS1. Silencing the S2→nwS1 pathway with inhibitory DREADDs disrupts locomotion, but silencing the M1→nwS1 pathway has little effect. Using *ex vivo* patch-clamp recording, we map the synaptic connectivity of long-range projections from S2 and M1 to nwS1. The S2→nwS1 pathway preferentially targets parvalbumin (PV) and somatostatin (SST)-expressing inhibitory interneurons, whereas the M1→nwS1 pathway connects to all neuronal subtypes. These results suggest that nwS1 and S2 coordinate paw movement during locomotion by engaging local inhibitory interneurons in nwS1.

## Materials and Methods

### Animals

All animal procedures conformed to NIH standards and were approved by the Institutional Animal Care & Use Committee at the University of Michigan. We report data from off-spring of B6.Cg-Gt(ROSA)26Sortm9(CAG-tdTomato)Hze/J mice (Jackson Laboratories, 007909) crossed to either B6;129P2-Pvalbtm1(cre)Arbr/J (Jackson Laboratories, 017320), Viptm1(cre)Zjh/J (Jackson Laboratories, 031628), or B6N.Cg-Ssttm2.1(cre)Zjh/J mice (Jackson Laboratories, 018973) on a mixed background. Mice were housed in a vivarium with a light–dark cycle of 12 h for each phase, and were aged 8–12 weeks at the start of experiments. Both sexes were used in all experiments. For behavioral testing, mice were matched by age and body weight.

### Stereotaxic injection

Mice were anesthetized with 1% isoflurane and mounted in a stereotaxic apparatus (David Kopf Instruments). Body temperature was maintained with a thermal blanket (Harvard Apparatus). The skull was exposed by a small incision on the scalp, cleaned with sterile saline and scored with a dental drill. A borosilicate pipette was beveled (30–50 μm internal diameter) and front-loaded with injection fluid. Injections were administered unilaterally via a craniotomy over the left hemisphere at a speed of 1 nl/s using the pipette mounted on an oil-based hydraulic micromanipulator (Narishige). For the mice used in the nwS1 silencing experiment, a headpost was attached to the skull using superglue (Krazy Glue^TM^) to facilitate sequential injections of saline and muscimol into the same mouse. For other experiments, the incision was closed using glue (Vet Bond, 3M).

### Rotarod assay

To assess motor coordination, we used the accelerating rotarod assay (Jones and Roberts, 1968). Mice performed two testing sessions separated by a 3-hour interval. Each testing session consisted of three trials, resulting in six trials per day for a given mouse. Mice were placed on the rotarod, and the speed was increased from 0 to 40.2 rpm over 300 seconds. Mice were rested for 5 minutes between trials in each session. For each trial, time spent on the rod was recorded. Trials were stopped when the mouse either fell off the rod or clung to the rod for three consecutive turns. The experimenter was blind to treatment conditions except when the effects of different treatments were compared in the same mouse.

### Locomotion assay

Mice were water-restricted for 1 week (1 ml per day) and trained to walk on the custom-built transparent runway (3 sessions, 1 session / day) by providing water in a dish hung at the end of the runway. The transparent runway was 40 cm long, 10 cm wide and 7.5 cm tall and elevated 30 cm from the floor. A training session consisted of five ‘runs’ on the runway. At the end of each run, mice were allowed to drink up to 0.2 ml of water from the dish which was refilled manually by the experimenter. Mice typically walked straight to the dish after training for 3 consecutive days. On the fourth day, all mice were subjected to two test sessions (five runs per session) on the runway. The first test session was preceded by a saline injection, and the second session by a muscimol injection. For each run, 0.1 ml of water was provided so that the total consumption remained the same as the training day (1 ml). Paw movements were recorded at 120 Hz with two GoPro cameras placed 25 cm from the right panel and 20 cm from the rear end of the runway.

### nwS1 silencing experiments

After recovery from headpost surgery (7–10 days), mice were water-restricted for 1 week (1 ml per day) and trained to walk on the custom-built transparent runway (3 days) by placing water at the end of the runway. Each mouse received a saline injection and a muscimol injection. nwS1 (7 mice; 0.7 mm posterior, 2.5 mm lateral from bregma) or wS1 (2 mice; 0.7 mm posterior, 3.7 mm lateral from bregma) was stereotaxically injected with 70nl saline. Injections occurred at three different depths (200, 400, 600 μm) from the pial surface. The craniotomy was sealed with silicone sealant (KwikCast^TM^) following injection, and covered with a thin layer of dental cement. Rotarod and locomotion assays were carried out after recovery from saline injection (1 h). Next, a fluorescently labeled muscimol-BODIPY conjugate (Hello Bio) was prepared in sterile saline solution (1 mg/ml) and 70 nl injected into nwS1 or wS1 at three different depths (200, 400, 600 μm) from the pial surface. Mice were euthanized after experiments and BODIPY fluorescence was examined.

### Grid shock test

Mice were placed in an acrylic chamber equipped with a stainless steel grid floor through which electric shock could be delivered (MazeEngineers). The shock generator delivered an output current pulse (2 s) starting at 0 mA every 20 sec and increasing with 0.1 mA up to a maximal 0.5 mA using a software provided by the manufacturer. Shock sensitivity threshold was determined as the current that elicited jump.

### Retrograde labeling experiments

Brain regions that project to nwS1 and wS1 were specifically analyzed using retrograde labelling. nwS1 was stereotaxically injected with 80-100 nl cholera toxin subunit B (CTB) conjugated to Alexa Fluor 488 (CTB-488, Invitrogen), and wS1 with CTB-Alexa Fluor 555 (CTB- 555, Invitrogen). The craniotomy was sealed with dental cement before closing the surgical site with glue. After 3 weeks, mice were perfused transcardially with ice-cold 0.9% sterile saline followed by 10% neutral-buffered formalin. Brains were embedded in agarose and fixed overnight (10-12 h) in 10% neutral-buffered formalin before coronal sectioning (80–100 μm).

After washing in PBS, brain sections were mounted on glass slides in DAPI-containing mounting medium (H-1200, Vector Laboratories). Images of processed tissues were acquired using a confocal microscope (Nikon A1) and analyzed using ImageJ. Distinct zones showing fluorescent labeling were selected as regions of interest (ROIs). Background fluorescence was determined by averaging intensity values over regions adjacent to the ROI. The background value was subtracted from the average fluorescence intensity of each ROI. Each ROI was assigned to a brain area according to the mouse brain atlas (Paxinos and Franklin, 2012). The background-adjusted fluorescence intensity was summed over each ROI and across brain sections to estimate the strength of retrograde labeling. Cortical and thalamic areas were analyzed separately. The sum of background-adjusted fluorescence signal in each brain area was divided by the grand sum of all areas in cortex or thalamus. Co-localization was quantified as Manders’ coefficients using ImageJ after setting a background threshold in the green and red channels separately.

### DREADD experiments

For pathway-specific inactivation, AAV9-hSyn-DIO-hM4D(Gi)-mCherry (titer 2.5 × 10^13^ GC/ml, Addgene 44362) and retrograde AAV encoding Cre recombinase (AAVretro-hSyn-HI-eGFP-Cre (titer 1.17 x 10^13^ GC/ml, Addgene 105540) were stereotaxically injected into S2 or M1 and nwS1, respectively (coordinates from bregma: S2, 1.2 mm posterior, 5.5 mm lateral; M1 1.0 mm anterior, 1.0 mm lateral). For cell type-specific inactivation, AAV9-hSyn-DIO-hM4D(Gi)-mCherry was injected into nwS1 of PV-Cre or SST-IRES-Cre mice. Injection was targeted to three different depths at 200, 400 and 600 μm from the pial surface (50-100 nl at each depth). The craniotomy was sealed with dental cement before closing the surgical site with glue. Three weeks after surgery, mice received an intraperitoneal injection of a water-soluble version of CNO (CNO-2HCl, 4.13 mg/kg, Tocris) or saline and underwent behavioral testing 1 h later. CNO injections were administered once a day before the first trial. Mice were euthanized after experiments and hM4Di expression was examined using mCherry signal.

### Neuronal ablation

Diphtheria toxin receptor (DTR) was expressed in nwS1 by stereotaxic injection of AAV1-Flex DTR-IRES-GFP (titer 2.93 × 10^13^ GC/ml, UM vector core) into PV-Cre; td-Tomato^fl/+^ mice (coordinates from bregma: posterior, 0.7 mm; lateral, 2.5 mm). The plasmid for AAV1-flex-DTR-GFP was a generous gift from Eiman Azim (Salk). Injection occurred at depths of 200, 400, 600 μm (40 nl per depth). At least 2 weeks after surgery, mice received one intraperitoneal injection of diphtheria toxin (DTX, diluted to 20 ng/µl with saline, 30 ng/g) or saline on 2 consecutive days. Mice were subjected to the rotarod test after 7 days following DTX/saline injection. Mice were then euthanized, and the loss of PV neurons was examined by comparing tdTomato signal between the two hemispheres.

### *Ex vivo* axonal fluorescence imaging and optogenetic stimulation

AAV5-CamKIIa-hChR2 (H134R)-EYFP (titer: 1.5 × 10^13^ GC/ml, Addgene, 26969) was stereotaxically injected into S2 or M1 in mice aged 4-6 weeks. Injection was targeted to three different depths at 200, 400 and 600 μm from the pial surface (50-100 nl at each depth). After 2- 4 weeks, mice underwent acute transcardial perfusion with ice-cold, aerated dissection solution. Acute coronal slices (300 μm thick) containing nwS1 and the S1 barrel field were prepared in a semi-frozen 300 mOsm dissection solution containing (in mM): 93 choline chloride, 2.5 KCl, 1.25 Na_2_H_2_PO_4_, 30 NaHCO_3_, 25 D-glucose, 20 HEPES, 3 Na pyruvate, 5 Na ascorbate, 10 MgCl_2_, and 0.5 CaCl_2_. Final osmolarity was adjusted with sucrose to 300–310 mOsm. The solution was continually perfused with 95% O_2_ and 5% CO_2_ prior to and during the slicing procedure. Slices were transferred to fresh dissection solution, warmed to 32°C to recover for 15 minutes, then transferred to 300–310 mOsm normal artificial cerebrospinal fluid (ACSF) containing, in mM: 125 NaCl, 2.5 KCl, 1.25 Na_2_H_2_PO_4_, 25 NaHCO_3_, 10 D-glucose, 1 kynurenate, 4 MgCl_2_, and 4 CaCl_2_, to recover at room temperature for at least 60 minutes prior to recording. eYFP and tdTomato signals were visualized using 470 nm and 565 nm LEDs respectively (Cool LED) delivered through a 40x upright microscope objective (Olympus). Fluorescence was detected using a CCD camera and visualized using Ocular software (Scientifica). To analyze spatial distribution of axonal fluorescence in nwS1 of brain slices expressing ChR2, a 250 mm x 800 mm ROI was manually selected. The ROI was selected based on the fluorescence distribution along the axis perpendicular to the pial surface. Fluorescence intensity was extracted using ‘Plot Profile’ function ImageJ. For photostimulation, blue light was delivered to the brain slice through a 470 nm LED (pE-100, CoolLED) coupled to 40 x upright microscope objective (Olympus). The light density was estimated to be ∼3.82 mW/mm^2^ at the level of the specimen. Each sweep consisted of two 5 ms pulses with a 200 ms inter-pulse interval, and each cell received five sweeps with a 45 s inter-sweep interval.

### Electrophysiology

Neurons were visualized with infrared-DIC optics. Whole-cell voltage clamp recordings were conducted using glass microelectrodes (borosilicate glass, pipette resistance 4–6 MΩ). Signals were acquired using the Multiclamp 700B amplifier (Molecular Devices) and pClamp 10.7, and digitized with Digidata 1550B. Signals were sampled at 10 kHz, low-pass filtered to 2.5 kHz and analyzed using Graphpad Prism software. Inhibitory interneuron subtypes were identified by tdTomato expression and pyramidal neurons by their morphology and lack of tdTomato fluorescence marker. The laminar location of neurons was determined by distance from the pia. Extracellular normal ACSF solution (described above) was continually aerated with 95% O_2_ and 5% CO_2_ and recirculated throughout recording. Temperature was constantly monitored and kept at 32°C by an in-line feedback-controlled heater. Unless stated otherwise, electrophysiological experiments were performed in voltage-clamp mode at −60mV using an internal solution (containing, in mM: 120 K-gluconate, 5 NaCl, 10 HEPES, 1.1 EGTA, 4 MgATP, 0.4 Na_2_GTP, 15 phosphocreatine, 2 MgCl_2_, 0.1 CaCl_2_) at 290–295 mOsm. Alexa 488 (Invitrogen) was mixed into the intracellular solution and used to visualize the morphology of recorded neurons. TTX (1 µM) and 4-AP (100 µM) were added to ACSF unless mentioned otherwise. We excluded recordings with a series resistance of >25 MΩ at the beginning or >30 MΩ at the end of the recording. All data were acquired and analyzed using custom Clampex software.

### Electrophysiology: controlling for hChR2 expression variability

We used a camera attached to the electrophysiology rig to acquire an image of hChR2-EYFP fluorescence at the recording site in nwS1. EYFP fluorescence at the recording site was normalized to autofluorescence (Fig. 1 inset), and this ratio was used as a proxy for ChR2 expression in each slice. We found that the mean amplitude of eEPSCs of excitatory cells in each slice had a strong positive linear relationship with hChR2-EYFP fluorescence (R = 0.80, p<0.001) (Fig. 1). To test if this relationship is linear regardless of the injection site, we compared R values in recordings obtained from slices after S2 or M1-targeted injection.

**Figure 1.**
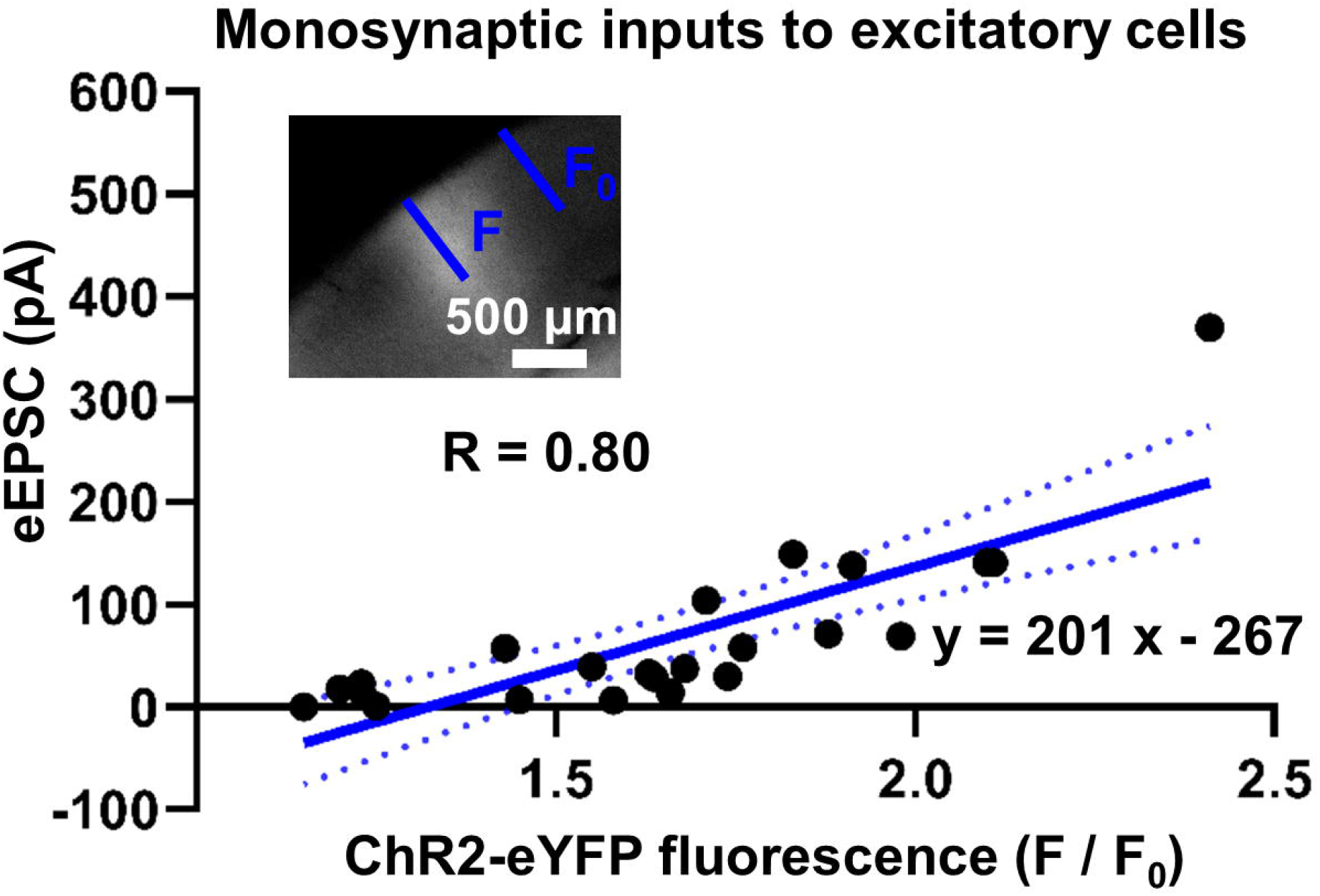
The level of hChR2-eYFP expression in long-range cortical projections is linearly related to light-evoked synaptic currents in excitatory cells of the target area. EYFP fluorescence (F) at the recording site was normalized to autofluorescence (F_0_) and plotted against the average eEPSC of excitatory cells in each slice for all experiments (R = 0.80, p<0.001; 23 slices). The mean amplitude of eEPSC of excitatory cells in each slice has a strong positive linear relationship with hChR2-EYFP fluorescence (R = 0.80, p<0.001). The line of best fit (solid blue) and 95% confidence interval (dashed blue) are indicated.

Comparisons of EYFP fluorescence or excitatory cell eEPSC amplitude revealed no statistical difference between S2 and M1 experiments (Mann-Whitney test, p > 0.1 for each comparison). The amplitude of excitatory cell eEPSCs strongly correlated with hChR2-EYFP expression regardless of the injection site (S2: R = 0.75; M1: R = 0.84). Therefore, eEPSC amplitude in excitatory cells can be used to normalize variable ChR2 expression when synaptic strengths are compared across different inhibitory subtypes, a standard practice in the field. We determined the line of best fit and the 95% confidence interval (CI) that describes the relationship between eEPSC amplitude and ChR2 expression. To compare eEPSC amplitudes across different neuronal subtypes after adjusting for variability in hChR2 expression, we asked if the eEPSC amplitude values of each subtype are significantly above or below 95% CI of this linear model.

### Statistics

The data were analyzed using MATLAB, Prizm software and ImageJ. For the rotarod data, the latency-to-fall values of individual mice are reported as means and SEMs. Rotarod performance was compared between groups by using a paired or unpaired *t*-test depending on whether a treatment was made in the same or different animals. Hind paw angles are reported as mean and standard error of the mean (SEM), and compared between mice that received saline and those that received muscimol or CNO using a paired *t*-test. eEPSC amplitudes of individual pairs of inhibitory and excitatory neurons are reported as means and SEMs. The Mann-Whitney test was used to compute the statistical significance of different eEPSC amplitudes among different neuronal subtypes. The degree to which S2 or M1 input was biased toward an inhibitory neuron over the nearby excitatory neuron was quantified by calculating input bias for each pair using the following formula: (difference in eEPSC amplitudes of inhibitory and nearby excitatory neurons) / (sum of eEPSC amplitudes of inhibitory and nearby excitatory neurons).

The Wilcoxon signed-rank test was used to test the null hypothesis that the mean input bias is zero. For all statistical tests, we used a significance criteria of p < 0.05. Probability values < 0.001 are described as p < 0.001. We did not use statistical methods to predetermine sample sizes, but our sample sizes are consistent with those reported in the field. All statistical tests were two-sided. Effect size is reported as a Hedges’ *g* value.

### Data and code availability

The datasets generated in this study and the code used for their analysis are available from the corresponding author upon request.

## Results

### Silencing nwS1 disrupts hind paw movement during locomotion

S1 stimulation is known to be sufficient to induce movements like whisker retraction and locomotion in mice (Matyas et al., 2010, Karadimas et al., 2020). A recent study demonstrated that silencing S1 leads to a decreased number of locomotion bouts in open field tests, indicating that S1 plays an important role in locomotion (Karadimas et al., 2020). However, specific aspects of locomotion disrupted by S1 perturbation were not characterized. If S1 is important for encoding somatosensory and/or proprioceptive feedback during locomotion, as suggested by previous studies in rodents (Chapin and Woodward, 1982b), then silencing nwS1 is expected to degrade precise paw placement during walking, thereby disrupting efficient locomotion (Akay et al., 2014). We tested this possibility using the accelerating rotarod and runway locomotion assays.

nwS1 was specifically silenced by stereotactic injection of the BODIPY-conjugated muscimol, a GABA-A receptor agonist. *Post hoc* histology revealed that the muscimol injection sites were located around the following bregma coordinates: lateral, 2.03 ± 0.145 mm; posterior, 0.83 ± 0.135 mm; these include the S1 hind limb, shoulder and dysgranular areas. The injected sites did not overlap with whisker S1 or nearby M1 areas (Fig. 2*A*) (Paxinos and Franklin, 2012). Each mouse received a sham injection with saline solution and performed the rotarod test. BODIPY-muscimol was injected into nwS1 (7 mice) or whisker S1 (2 mice), and performance was assessed again. Hind paw movements during the rotarod test were recorded using a camera (30 Hz) (Fig. 2*B*, *C*). Muscimol injection into nwS1 impaired rotarod performance and led to a significantly shorter ‘latency to fall’ time compared to saline injection in the same mice (‘S’, saline: 107 ± 7.01 s; ‘M’, muscimol: 61.4 ± 7.26 s; 7 mice; t(6) = 5.11, p = 0.0022, g = 2.40, paired *t*-test; Fig. 2*D*). This effect was specific to nwS1; muscimol injection into wS1 had little effect on rotarod performance (saline: 115 ± 23.3 s; muscimol: 136 ± 12.0 s; 2 mice; Fig. 2*D*).

**Figure 2.**
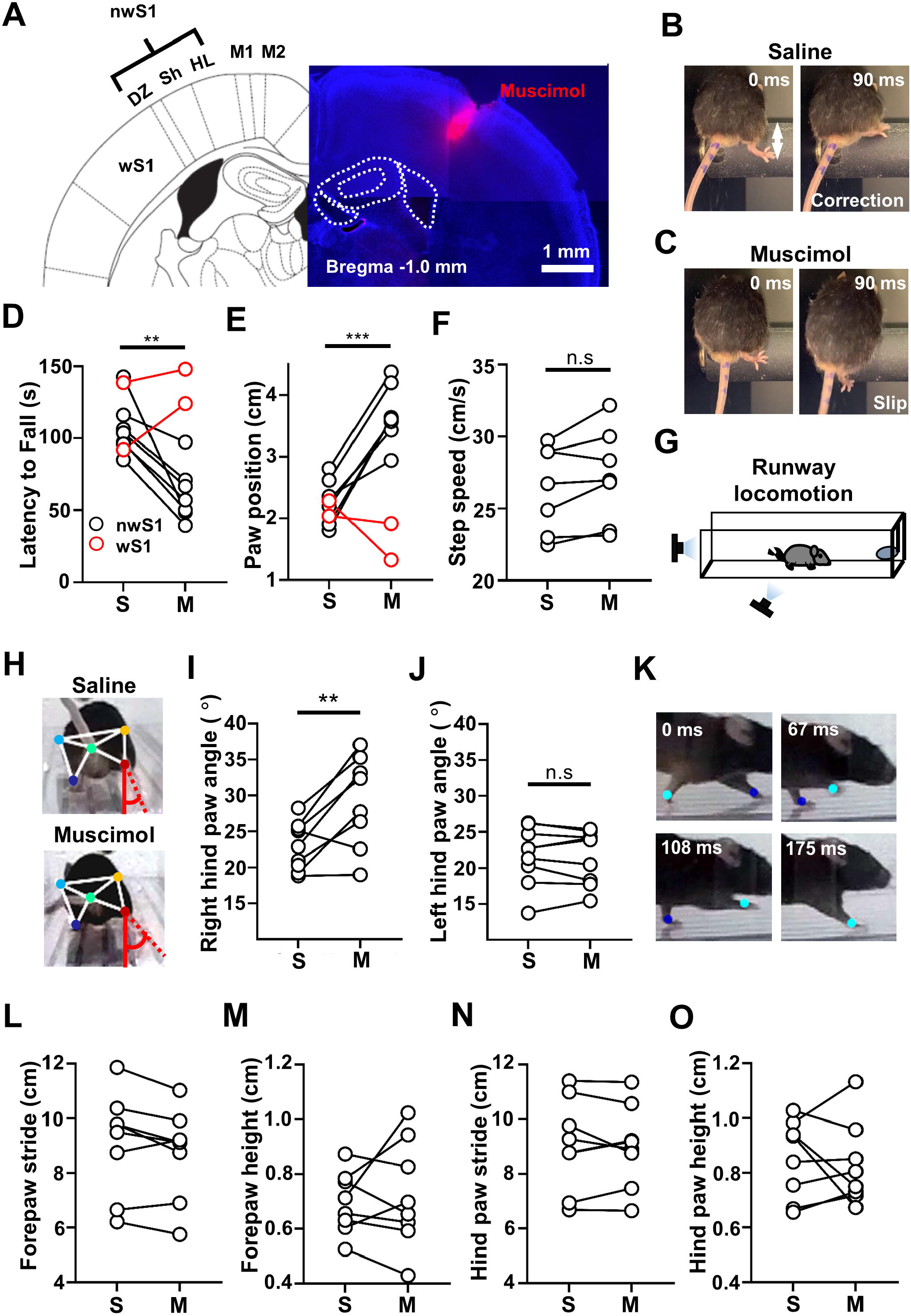
Silencing nwS1 disrupts movements of the hind paw during locomotion. **(A)** Representative image of a coronal brain section showing muscimol injection localized to nwS1. DZ: dysgranular zone; Sh: shoulder area; HL: hind limb area; M1: primary motor cortex; M2: secondary motor cortex. **(B, C)** Images showing hind paw placement during the rotarod performance after (B) saline (‘S’) or (C) muscimol (‘M’) injection into nwS1. The distance from the right hind paw of the mouse to the apex of the rod (white arrow) at the initiation of the corrective step was measured. After muscimol injection, the paw frequently slipped off the rod (C). **(D)** Average rotarod performance of mice measured as the latency to fall before and after muscimol inactivation of nwS1 or wS1. Each mouse performed a set of three trials after saline injection and another set of three trials after muscimol injection with a 3-hour break in between. The latency to fall was averaged across three trials for each condition. Mean ± SEM for nwS1 injection: saline, 107 ± 7.00 s (7 mice); muscimol, 61.4 ± 7.26 s (7 mice); ** p = 0.0022, paired *t*-test. Mean ± SEM for wS1 injection: saline, 115 ± 23.3 s (2 mice); muscimol, 136 ± 12.0 s (2 mice). **(E)** Average paw position at the time of initiation of the corrective step before and after muscimol inactivation of nwS1 or wS1. Mean ± SEM for nwS1 injection: saline, 2.28 ± 0.136 cm (7 mice); muscimol, 3.68 ± 0.181 cm (7 mice); *** p < 0.001, paired *t*-test. Mean ± SEM for wS1 injection: saline, 2.16 ± 0.124 cm (2 mice); muscimol, 1.62 ± 0.294 cm (2 mice). **(F)** Average speed of the corrective step before and after muscimol inactivation of nwS1. Mean ± SEM for nwS1 injection: saline, 26.4 ± 1.130 cm/s (7 mice); muscimol, 27.3 ± 1.241 cm/s (7 mice); p = 0.072, paired *t*-test. **(G)** Experimental set-up for the locomotion assay. Mice were water-restricted and trained to walk on the transparent runway to drink water from a dish hung at the end of the runway. **(H)** Rear view images of a mouse with saline or muscimol injection into nwS1. Example labels used for DeepLabCut analysis of a movie showing a mouse walking on the runway. Dark blue, left hind paw; Red, right hind paw; Light blue, left knee; Orange, right knee; Jade, bottom of tail. Hind paw angle was measured between the vertical axis (solid red) and the midline along the length of the hind paw (dashed red). **(I, J)** Right (I) and left (J) hind paw angles in mice after injection of saline or muscimol into nwS1. Each data point represents an average of the hind paw angles during multiple strides observed in a single mouse. (I) Saline: 22.95° ± 1.10; muscimol: 30.43° ± 2.34; n = 9 mice, ** p = 0.009, paired *t*-test. (J) Saline: 21.77° ± 1.35; muscimol: 21.62° ± 1.24; p = 0.737, n = 9 mice, p = 0.737, paired *t*-test. **(K)** Side view images of forepaws during a stride on the runway. Example labels used for DeepLabCut analysis of a movie showing a mouse walking on the runway. Dark blue, left forepaw; Light blue, right forepaw. **(L, M)** Forepaw stride length (*L*) and height (*M*) in mice after injection of saline or muscimol into nwS1. Each data point represents an average of the stride length or height measured during multiple strides observed in a single mouse. (L) Saline: 9.11 cm ± 0.67; muscimol: 8.73 ± 0.59; n = 8 mice, p = 0.069, paired *t*-test. (M) Saline: 0.69 cm ± 0.67; muscimol: 0.72 ± 0.67; n = 8 mice, p = 0.597, paired *t*-test. **(N, O)** Hind paw stride length (*N*) and height (*O*). (N) Saline: 9.08 cm ± 0.59; muscimol: 9.02 ± 0.53; n = 8 mice, p = 0.763, paired *t*-test. (O) Saline: 0.85 cm ± 0.05; muscimol: 0.83 ± 0.05; n = 8 mice, p = 0.654, paired *t*-test.

We noticed that the corrective step of the right hind paw was severely impaired following the muscimol injection into left nwS1 (Fig. 2*B*, *C*). To quantify the deficit in paw movement, we measured the distance travelled by the right hind paw from the apex of the rotating rod before the corrective step occurred (Cao et al., 2015). Compared to the saline injection in the same mouse, muscimol injection into nwS1 significantly increased the apex-to-paw distance such that mice rarely initiated the corrective step before the paw slipped off the rod (saline: 2.28 ± 0.136 cm; muscimol: 3.68 ± 0.181 cm; 7 mice; t(6) = 8.76, p<0.001, g = 3.30, paired *t*-test; Fig. 2*E*).

Therefore, it took a larger displacement of the right hind paw along the rotating rod before the animal initiated the corrective step to maintain balance. In contrast, silencing wS1 had little effect (saline: 2.43 ± 0.197 cm; muscimol: 1.82 ± 0.331 cm; 2 mice). The delayed corrective step may be caused by impairments in representing somatosensory input such as the paw / hip position, in generating a motor command for the step or in using somatosensory information to guide the step. To gain insight into the nature of the deficit caused by S1 inactivation, we compared the speed of corrective step before and after nwS1 silencing in the same mouse to test whether the motor command is affected. The average speed of corrective step was not significantly different between saline and muscimol treatments in the same mouse (saline: 26.4 ± 1.130 cm/s; muscimol: 27.3 ± 1.241 cm/s; 7 mice; t(6) = 2.18, p = 0.072, g = 0.28, paired *t*- test; Fig. 2*F*). Therefore, while the corrective step was initiated at a position further away from the apex after nwS1 silencing, the movement itself was normal once it was initiated. Our results therefore suggest that nwS1 is specifically and critically involved in sensing the paw / hip position during locomotion on the rotarod, building on a classic study in spinal cats (Grillner and Rossignol, 1978).

We also tested whether nwS1 is important for over-ground locomotion in a separate group of mice. Water-restricted mice were trained to walk on a transparent runway to collect water (Fig. 2*G*). After three daily training sessions, all the mice went straight to the water when placed on the runway. On the fourth day, mice received a saline injection into nwS1, and their performance was tested on the runway. Each mouse performed five trials and was allowed to collect up to 0.1 mL of water per trial. After resting for 1 h, the same mouse then received a unilateral injection of BODIPY-conjugated muscimol into the left nwS1 and was tested on the runway again. Paw movements during locomotion were recorded using cameras placed at two different angles to acquire rear and side views (120 Hz). Hind paw positions were extracted from the recorded movies using the machine learning-based platform DeepLabCut (Mathis et al., 2018). We matched the rear and side view images frame by frame to identify the onset of swing phase in each stride during locomotion. The angle of the contralateral (right) hind paw at the onset of the swing phase was significantly greater after nwS1 silencing with muscimol (saline: 22.95° ± 1.10; muscimol: 30.43° ± 2.34, n = 9 mice; t(8) = 3.45, p = 0.009, g = 1.36, paired *t*-test; Fig. 2*H*, *I*; Movie 1). In contrast the ipsilateral (left) hind paw angle was unaffected by nwS1 inactivation (saline: 21.77° ± 1.35; muscimol: 21.62° ± 1.24; n = 9 mice; t(8) = 0.348, p = 0.737, g = 0.04, paired *t*-test) (Fig. 2*J*). Silencing nwS1 disrupted the trajectory of locomotion such that the body tended to tilt towards the left during walking. This demonstrates that nwS1 inactivation disrupts the orientation of paw movements during locomotion. In contrast, the locomotion speed was not affected by nwS1 silencing (saline: 25.27 cm/s ± 3.62; muscimol: 24.38 cm/s ± 3.39; t(8) = 0.502, p = 0.631, g = 0.09, paired *t*-test), indicating that gross locomotion was not altered.

nwS1 inactivation could have a broad impact on limb kinematics during locomotion in addition to the altered paw orientation. To test this possibility, we measured average stride length and paw height for right forepaw and right hind paw using side-view images (Fig. 2*K*). Stride length was measured as the distance between two successive placements of the right forepaw. Paw height was measured as the vertical distance between the paw at the highest position during a stride and the runway floor. We did not observe significant difference in the stride length (Saline: 9.11 cm ± 0.67; muscimol: 8.73 ± 0.59; n = 8 mice; t(7) = 2.15, p = 0.069, g = 0.217, paired *t*-test) or paw height (Saline: 0.69 cm ± 0.67; muscimol: 0.72 ± 0.67; n = 8 mice; t(7) = 0.554, p = 0.597, g = 0.181, paired *t*-test) (Fig. 2*L, M*). Right hind paw also did not show significant differences in the stride length (saline: 9.08 cm ± 0.59; muscimol: 9.02 ± 0.53; n = 8 mice; t(7) = 0.313, p = 0.763, g = 0.026, paired *t*-test) or paw height (saline: 0.85 cm ± 0.05; muscimol: 0.83 ± 0.05; n = 8 mice; t(7) = 0.468, p = 0.654, g = 0.157, paired *t*-test) (Fig. 2*N, O*). Therefore, nwS1 inactivation primarily impacts the contralateral hind paw orientation rather than resulting in broadly impaired paw movement. Note that a comprehensive analysis of paw kinematics is needed to exclude possible effects on forepaws.

Impaired locomotion can result from changes in pain perception during nwS1 inactivation (Favorov et al., 2019). To test this possibility, we compared sensitivity to mild electric shock before and after muscimol injection into nwS1. Mice were placed in a chamber equipped with a shock grid. The shock threshold was defined as the lowest current required to elicit jump behavior. We saw a slight increase in average shock threshold after muscimol injection into nwS1 compared to saline injection into the same mouse. The observed change was not statistically significant (saline: 0.24 mA ± 0.024; muscimol: 0.30 mA ± 0.055; n = 5 mice; t(4) = 1.50, p = 0.208, g = 0.632, paired *t*-test). Based on these results, we conclude that nwS1 inactivation specifically disrupts hind paw orientation during over-ground locomotion.

### Identification of cortical regions that project to nwS1

S1 is reciprocally connected with other cortical areas, including M1, S2 and the posterior parietal cortex, which together form a cortical sensorimotor subnetwork (Zingg et al., 2014). Thalamocortical pathways, through which S1 receives sensory information ascending from periphery, are relatively well-documented. However, the role cortico-cortical connections in the regulation of locomotion remain unclear. Long-range cortical connections from M1 and S2 are thought to be topographically organized such that different subregions in S1 receive non-overlapping cortical inputs (Zingg et al., 2014, Minamisawa et al., 2018). However, cortical connections to subregions such as wS1 and nwS1 have rarely been compared in the same mouse brain and whether cortical inputs into nwS1 and wS1 are distinct was unclear. To address this issue, we injected the retrograde anatomical tracer CTB-Alexa 488 into nwS1 using stereotaxic coordinates taken from bregma (0.7 mm posterior, 2.5 mm lateral; n = 3 mice). In a subset of these mice, CTB-Alexa 555 was also injected into the wS1 (0.7 mm posterior, 3.7 mm lateral; n = 2 mice) (Fig. 3*A*). Retrogradely labeled nwS1-projecting neurons were distributed in several different cortical and thalamic areas. Note that fluorescent signals in S2 / ventral auditory area (AuV) were combined, as were ectorhinal cortex (Ect) / temporal association area (TeA), since there is no obvious anatomical border between these regions (Zingg et al., 2014).

**Figure 3.**
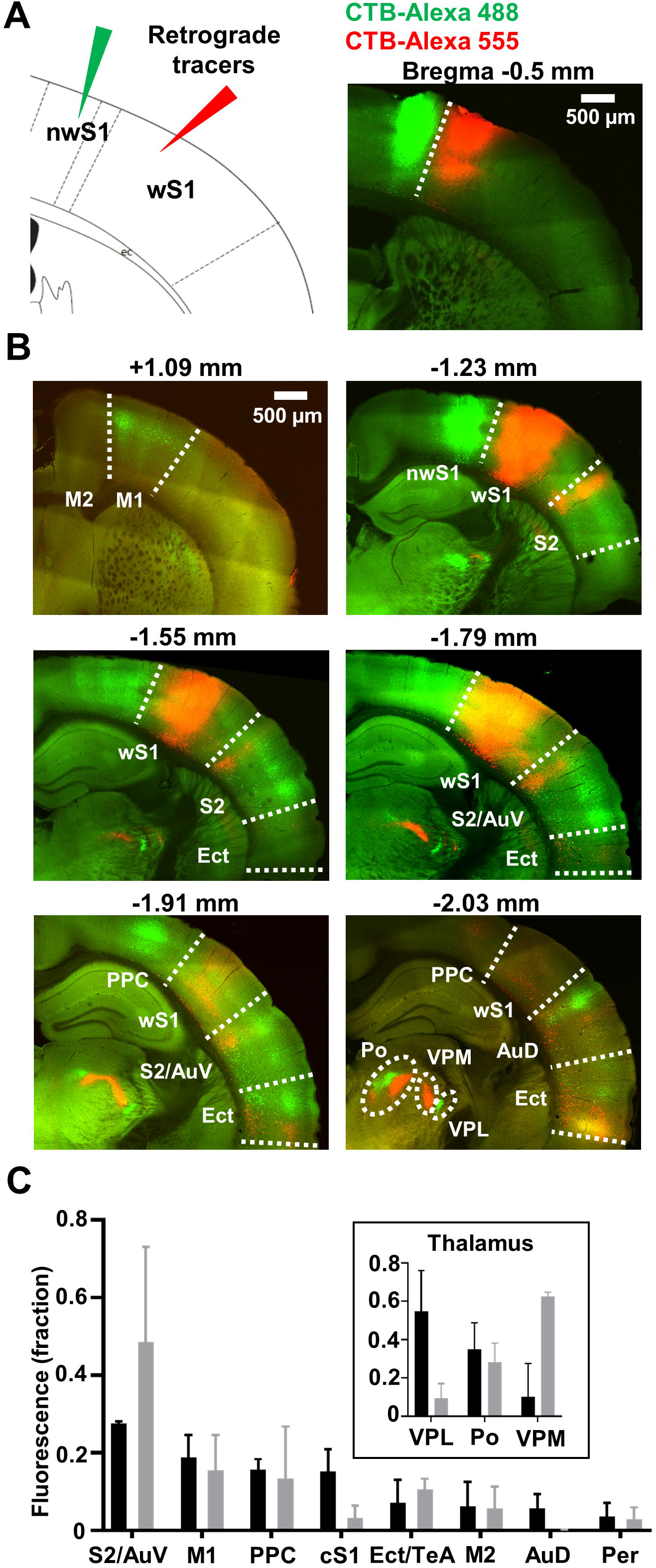
Tracing long-range anatomical inputs to nwS1. **(A)** Injection paradigm. Cholera toxin B conjugated with Alexa Fluor 488 (CTB-488) was injected into nwS1 (coordinates from bregma: posterior, 0.7 mm; lateral, 2.5 mm). Cholera toxin B conjugated with Alexa Fluor 555 (CTB-555) was injected into wS1 (posterior, 0.7 mm; lateral, 3.7 mm). **(B)** Representative images showing laminar distribution of labeling within S2 and M1 in coronal sections of mouse brains after retrograde tracer injections into nwS1 and wS1. AuD: dorsal auditory cortex; AuV: ventral auditory cortex; Ect: Ectorhinal cortex; M1: primary motor cortex; M2: secondary motor cortex; Po: posteromedial nucleus; PPC: posterior parietal cortex; S1: primary somatosensory cortex; S2: secondary somatosensory cortex; VPL: ventral posterolateral nucleus; VPM: ventral posteromedial nucleus. **(C)** Quantification of retrograde labeling based on the fraction of total fluorescence signal in cortical and thalamic (inset) areas. Mean ± SEM from n = 3 mice (nwS1; black) and n = 2 mice (wS1; gray); cS1: contralateral S1; TeA: temporal association area; Per: perirhinal cortex.

Cortical regions S2/AuV and M1 resulted in prominent labeling of nwS1-projecting neurons (Fig. 3*B*, *C*), consistent with previous studies in rats and squirrels (Cooke et al., 2012, Kim and Lee, 2013). Posterior parietal cortex (PPC), contralateral S1 (cS1), ectorhinal cortex (Ect) / temporal association area (TeA), secondary motor cortex (M2), dorsal auditory area (AuD) and perirhinal cortex (Per) also contained nwS1-projecting neurons in decreasing order (Fig. 3*C*). In thalamus, ventral posterolateral (VPL) and posteromedial nuclei (Po) nuclei showed strong labeling of nwS1-projecting neurons. Neurons projecting to wS1 were located in S2/AuV, M1 and Ect/TeA (Fig. 3*B, C*). Among thalamic nuclei, VPM and Po showed prominent labeling of wS1 projection neurons. To test whether nwS1- and wS1-projecting neurons form distinct groups, we quantified co-localization of Alexa 488 and Alexa 555 in the co-injected mice (n = 2 mice) using Mander’s colocalization coefficients (MCC). Two MCC values were calculated for cortex and thalamus separately in each mouse; MCC1, the fraction of red fluorescence (Alexa 555; wS1 injection) in compartments containing green fluorescence (Alexa 488; nwS1 injection) and MCC2, the fraction of green in compartments containing red. In cortical areas excluding injection sites, MCC1 and MCC2 were 0.087 ± 0.069 and 0.043 ± 0.044 (Mean ± STD; n = 2 mice). In thalamus, MCC1 and MCC2 were 0.006 ± 0.008 and 0.010 ± 0.010 (Mean ± STD; n = 2 mice). The low degree of overlap between wS1 and nwS1-projecting subregions was not due to difference in efficiency of retrograde labeling since co-injection of a mixture containing the two tracers into wS1 resulted in a high degree of overlap in both cortex and thalamus (cortex: MCC1 = 0.559, MCC2 = 0.672; thalamus: MCC1 = 0.671, MCC2 = 0.791; n = 1 mouse). Therefore, despite their proximity, nwS1 and wS1 receive distinct long-range cortical inputs and participate in distinct sensorimotor subnetworks.

Previous work suggests that S2 and M1 are reciprocally connected with nwS1 and form a lower limb/trunk sensorimotor subnetwork that is anatomically distinct from the whisker subnetwork (Zingg et al., 2014). We therefore asked if S2 and M1 provide inputs to nwS1 and form distinct streams of information by characterizing their projection patterns in nwS1. To compare distributions of S2 and M1 axons in nwS1, we performed anterograde tracing. We injected AAV5-CamKIIa-hChR2 (H134R)-EYFP into S2 and M1 (Fig. 4*A*). Injections were focused on S2 and M1 subregions that contained the most nwS1-projecting neurons (S2: 1.2 mm posterior, 5.5 mm lateral; M1: 1 mm anterior, 1 mm lateral). To compare cortical layers innervated by M1 versus S2 projections in nwS1, we quantified the spatial distribution of fluorescence from anterogradely labeled axons based on the distance from the pial surface. S2→nwS1 axons were concentrated in layer 2/3 and upper layer 4 (Minamisawa et al., 2018), whereas M1→nwS1 axons innervated layers 1, 2 and 6 (Fig. 4*B*) (Naskar et al., 2021). Our results show that S2 and M1 inputs to nwS1 form anatomically distinct cortico-cortical pathways.

**Figure 4.**
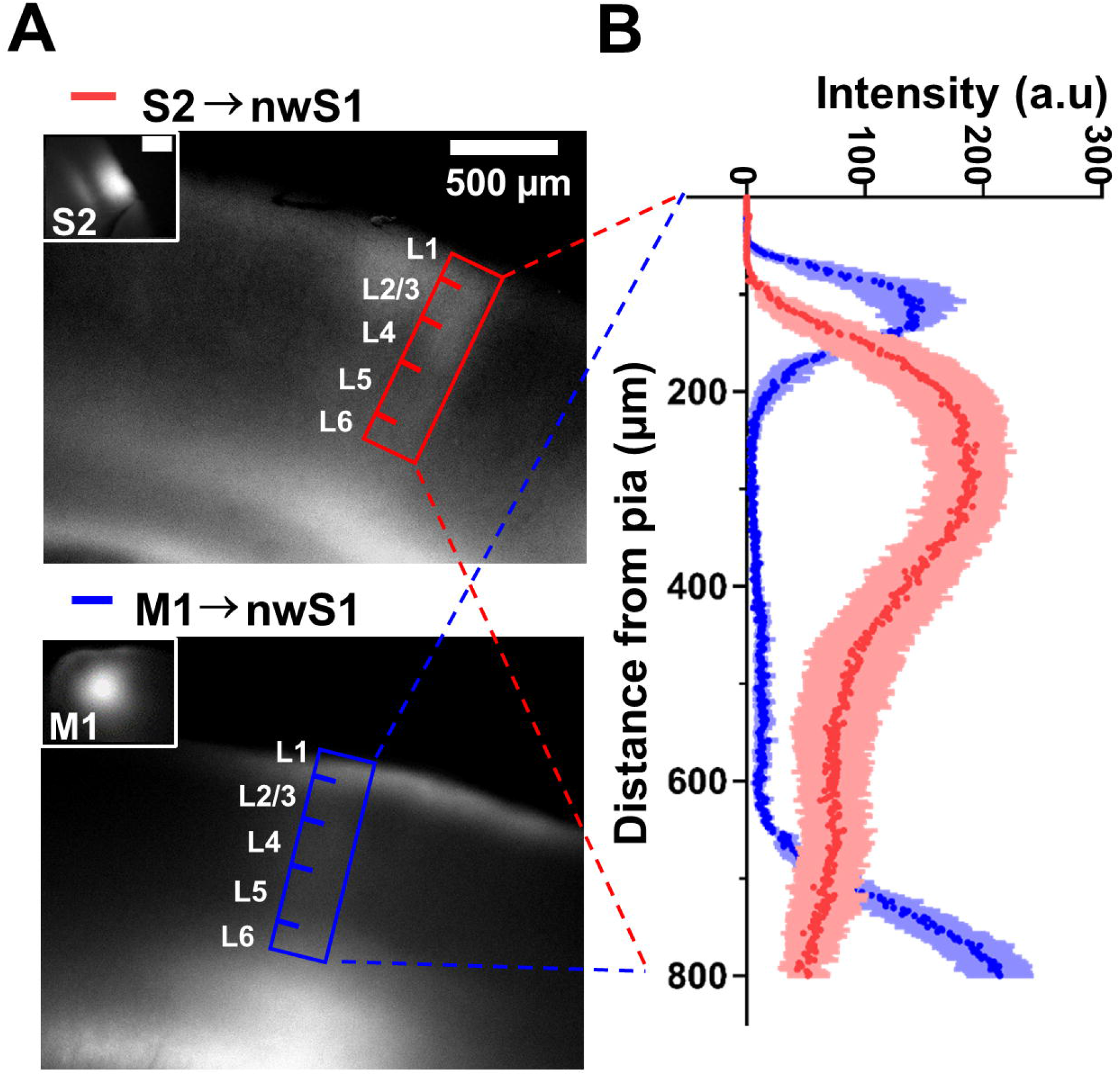
Distribution of S2 and M1 projections in nwS1. **(A)** Representative images showing S2 and M1 projections labeled with ChR2-eYFP. Images were taken from acute slices used in electrophysiological recording. Slices were isolated from brains injected with ChR2-YFP into S2 (upper panel) or M1 (lower panel). Insets show representative images of injection sites. Rectangles indicate the area in nwS1 selected for intensity analysis. Fluorescent signal in nwS1 indicates ChR2-expressing projections from S2 (red rectangle) or M1 (blue rectangle). **(B)** Laminar profile of ChR2-eYFP-labeled axons in nwS1. Each symbol represents the mean pixel intensity at the given distance from pia from 5 slices (4 mice) for each pathway. Shading indicates standard error of the mean across 5 slices (4 mice).

### S2→nwS1 input modulates motor performance

Having identified S2 and M1 as major cortical inputs to nwS1, we next asked whether the cortico-cortical connections formed by these inputs are involved in motor performance or not. Although mouse S1 plays an important role in efficient locomotion, as shown above and in other studies (Karadimas et al., 2020), the role of cortico-cortical projections was not explored. We selectively silenced the S2→nwS1 or M1→nwS1 pathway by expressing hM4Di, an inhibitory ‘designer receptor exclusively activated by designer drug’ (DREADD), using an intersectional approach (Armbruster et al., 2007). AAV encoding Cre-dependent hM4Di-mCherry was injected into S2 or M1, and retrograde AAV encoding Cre was injected into nwS1 (Fig. 5*A*). *Post hoc* histology showed that the expression of hM4Di-mCherry was localized to S2/AUD or M1 (Fig. 5*B*, *C*). Three weeks following hM4Di expression, mice received an i.p. injection of the DREADD agonist clozapine N-oxide (CNO-2HCl, 4.13 mg/kg) or saline and were tested on the rotarod. The CNO-injected group with S2→nwS1 silencing showed impaired rotarod performance, with a significantly shorter average ‘latency to fall’ time than the saline-injected group (saline: 222 ± 5.77 s, 7 mice; CNO: 183 ± 5.45 s, 8 mice; t(13) = 4.85, p < 0.001, g = 2.54, *t*-test; Fig. 5*D*). On the other hand, silencing the M1→nwS1 pathway did not affect mouse rotarod performance; no difference was observed between rotarod performance in the CNO and saline-treated groups in these mice (saline: 214 ± 19.1 s, 6 mice; CNO: 198 ± 12.3 s, 6 mice; t(10) = 0.665, p = 0.521, g = 0.38, *t*-test; Fig. 5*E*). Therefore, specific inactivation of S2→nwS1 projections degraded performance on the rotarod performance. This result also shows that CNO alone does not influence performance in the rotarod assay.

**Figure 5.**
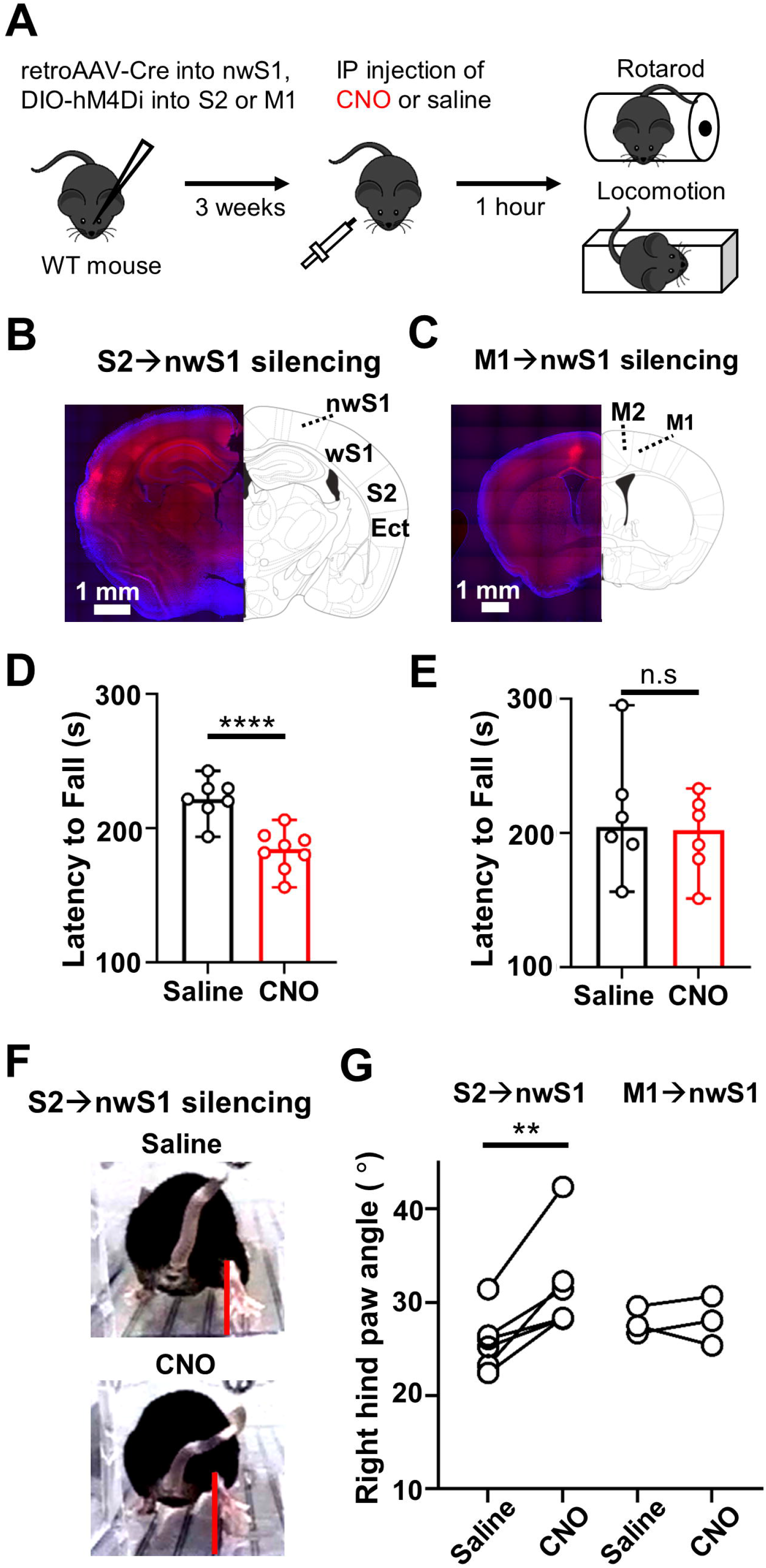
Silencing the S2→nwS1 pathway impairs both rotarod performance and runway locomotion. **(A)** Experimental paradigm. Retrograde AAV-Cre was injected into nwS1. AAV encoding the Cre-dependent hM4Di-mCherry was injected into S2 or M1. Three weeks later, mice received an intraperitoneal injection of CNO (CNO-2HCl, 4.13 mg/kg) or the equivalent volume of saline 1 h prior to testing on the rotarod and the locomotion assay. **(B, C)** Intersectional Cre-lox approach resulted in hM4Di-mCherry expression localized to S2 or M1. Demarcations are based on a mouse brain atlas (Paxinos and Franklin, 2012). **(D)** Average rotarod performance of mice expressing hM4Di in S2→nwS1 with CNO or saline injection. Saline: 222 ± 5.77 s, 7 mice; CNO: 183 ± 5.45 s, 8 mice; p=0.0003, *t*-test. **(E)** Average rotarod performance of mice expressing hM4Di in M1→nwS1 with CNO or saline injection. Saline: 214 ± 19.1 s, 6 mice; CNO: 198 ± 12.3 s, 6 mice; p=0.52, *t*-test. **(F)** Rear view images of a mouse walking on the runway before and after S2→nwS1 inactivation with hM4Di. **(G)** Mean right hind paw angles in mice before and after the S2→nwS1 or M1→nwS1 inactivation. Each data point represents a mean of the hind paw angles during multiple strides observed in a single mouse. S2→nwS1 inactivation; saline: 25.8 ± 1.30; CNO: 31.8 ± 2.24; 6 mice; p=0.008, paired *t*-test. M1→nwS1 inactivation; saline: 27.9 ± 1.50; CNO: 28.0 ± 2.64; 3 mice; p=0.930, paired *t*-test.

We also tested the effect of silencing cortico-cortical inputs on hind paw orientation during the runway locomotion assay in a separate cohort of mice. CNO injection into mice expressing hM4Di to silence S2→nwS1 projections increased the right hind paw angle at the onset of the swing phase of locomotion compared with saline injection in the same mice (saline: 25.8 ± 1.30; CNO: 31.8 ± 2.24; 6 mice; t(5) = 4.28, p = 0.008, g = 1.34, paired *t*-test; Fig. 5*F, G*). To test whether S2→nwS1 inactivation alters paw orientation as effectively as muscimol injection into nwS1 (Figs. 2, 5), we normalized paw angle following CNO or muscimol injection relative to the baseline in each animal. There were no significant differences between increased hind paw angle in hM4Di-CNO and muscimol treated mice (nwS1 inactivation: 1.34 ± 0.10, 9 mice; S2→nwS1 inactivation: 1.23 ± 0.05, 6 mice; p = 0.46, g = 0.39, Mann-Whitney test). CNO injection did not alter the hind paw angle in mice expressing hM4Di that silenced the M1→nwS1 pathway (saline: 27.9 ± 1.50; CNO: 28.0 ± 2.64; 3 mice; t(2) = 0.100, p = 0.930, paired *t*-test, g = 0.05). Based on these results, we conclude that the S2→nwS1 pathway is critically involved in maintaining the orientation of hind paw for efficient locomotion.

### S2→nwS1 pathway preferentially innervates inhibitory interneurons

The distinct functional outcomes of silencing S2 versus M1 inputs to nwS1 suggest that each pathway targets distinct local circuit elements. To gain insights into computations mediated by different long-range projections, we characterized neuronal subtypes that make up functional connections in the S2→nwS1 and M1→nwS1 pathways. Inhibitory interneurons expressing parvalbumin (PV), somatostatin (SST) or vasoactive intestinal peptide (VIP) together account for ∼82% of all interneurons in the cortex (Rudy et al., 2011). We reasoned that we could identify the major postsynaptic target(s) of S2 and M1 inputs by surveying these three interneuron subtypes and putative excitatory neurons. To this end, we recorded synaptic currents from genetically defined neuronal subtypes while photostimulating S2→S1 axonal projections in brain slices. Channelrhodopsin-2 (ChR2) was expressed in S2 by injecting AAV5-CamKIIa-hChR2 (H134R)-EYFP. The axon terminals expressing ChR2 were photostimulated with pulses from a 470 nm LED delivered through the microscope objective. We recorded evoked excitatory postsynaptic currents (eEPSCs) from genetically identified interneuron subtypes and putative excitatory cells (ECs) in nwS1 (Fig. 6*A*). Experiments were conducted in reporter mice generated by crossing PV-Cre, SST-Cre or VIP-Cre mice with Cre-dependent tdTomato mice (PV-Cre; tdTomato^lsl/+^; SST-IRES-Cre; tdTomato^lsl/+^; VIP-IRES-Cre; tdTomato^lsl/+^; Fig. 6*B*). nwS1 was identified in brain slices as the region lacks barrels and has a specific location relative to wS1 (Fig. 6*A*). We isolated monosynaptic eEPSCs by recording in the presence of tetrodotoxin (TTX) and 4-aminopyridine (4-AP) (Petreanu et al., 2009). TTX blocks action potentials, and 4- AP augments local depolarization of photostimulated ChR2-expressing axons by blocking voltage-gated potassium channels. The fluorescent dye Alexa 488 was included in into the intracellular solution regions to facilitate visualization of impaled cells during recording (Fig. 6*C*). A subset of mice were subjected to neuronal recordings that were made sequentially from pairs of inhibitory and excitatory cells (ECs) located within 50 µm of each other. eEPSC recordings were targeted to layers that matched spatial distribution of relevant axonal fluorescence (S2 experiments: layers 2/3 and upper 4). We repeated the same experiment in M1→nwS1 pathway (see Fig. 7). A brief photostimulation (5 ms) evoked monosynaptic EPSCs in many but not all cells (Fig. 6*D*). The peak eEPSC amplitude for PV-, SST-, VIP-expressing interneurons and ECs was 185 ± 35.8 pA (26 cells), 119 ± 18.9 pA (18 cells), 37.9 ± 10.7 pA (12 cells) and 34.3 ± 7.63 pA (29 cells) respectively (data expressed as mean ± SEM, Fig. 6*E*). Consistent with a recent finding in the mouse S2→ whisker S1 pathway (Naskar et al., 2021), PV-expressing interneurons received significantly greater monosynaptic input from S2 than both VIP interneurons and ECs (PV-VIP: p < 0.001, g = 0.95; PV-EC: p < 0.001, g = 1.16; Mann-Whitney tests). SST-expressing interneurons also showed significantly greater monosynaptic eEPSCs compared with both VIP-expressing interneurons and excitatory cells (SST-VIP: p = 0.002, g = 1.21; SST-EC: p < 0.0001, g = 1.43; Mann-Whitney tests). VIP-expressing interneurons and ECs did not show a significant difference (p=0.64, g = 0.09, Mann-Whitney test). These results suggest that the S2 → nwS1 pathway provides inputs to PV-expressing and SST-expressing local interneurons.

**Figure 6.**
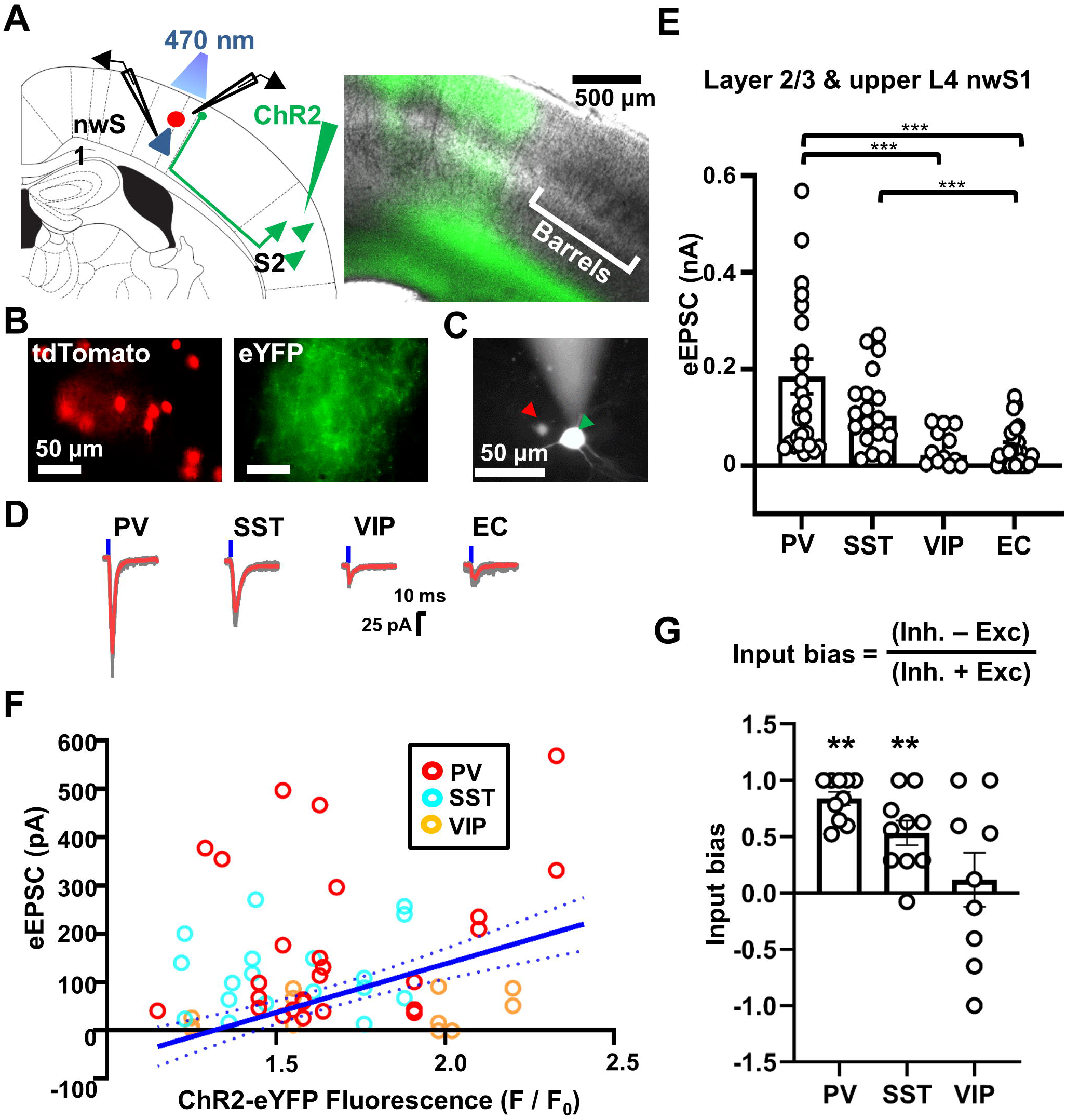
S2 projection preferentially innervates PV and SST-expressing interneurons in S1. **(A)** Left: Schematic of slice recording configuration. An interneuron (red) and a nearby putative excitatory neuron (blue) were recorded sequentially in pairs. See text for details. Right: Representative image of a slice fixed in 10% formalin after recording showing expression of ChR2-eYFP in S2 after stereotaxic injection of AAV5-CamKIIa-hChR2 (H134R)-EYFP into S2 (1.2 mm posterior, 5.5 mm lateral from bregma). **(B)** Left: Example neurons showing tdTomato expression. Right: Image from the same field of view showing the presence of ChR2-eYFP-expressing axons near the recording site. **(C)** Image showing two nearby cells that were sequentially recorded. Alexa-488 was mixed into intracellular solution for visualization. Red and green arrowheads indicate a tdTomato- expressing interneuron and a putative excitatory neuron, respectively. Putative excitatory neurons were determined by the lack of tdTomato signal. Images were acquired using a differential interference contrast microscope at ×4 magnification. **(D)** Example response traces from sequential paired recordings in slices expressing tdTomato in parvalbumin (PV)-, somatostatin (SST)-, or vasoactive intestinal peptide (VIP)-positive interneurons and excitatory cells (EC). Averages from five consecutive sweeps of optogenetically evoked excitatory postsynaptic currents (eEPSC, gray) are shown in red. Blue vertical lines indicate photostimulation (470 nm LED, 5 ms) delivered through the objective. Responses were acquired in the presence of tetrodotoxin and 4-aminopyridine. **(E)** eEPSC amplitudes of individual neurons in response to photostimulation of the S2→nwS1 projection. Each dot represents an average of five sweeps. Bar indicates mean; error bars indicate SEM. Mean ± SEM of peak eEPSC amplitude for PV-, SST-, VIP-expressing interneurons and ECs were 185 ± 35.8 pA (26 cells), 119 ± 18.9 pA (18 cells), 37.9 ± 10.7 pA (12 cells) and 34.3 ± 7.63 pA (29 cells) respectively. PV-expressing interneurons showed significantly greater monosynaptic eEPSCs than both VIP interneurons and excitatory cells (Mann-Whitney test, p<0.001 for each comparison). SST-expressing interneurons also showed significantly greater monosynaptic eEPSCs compared with both VIP-expressing interneurons and excitatory cells (Mann-Whitney test, p=0.002 for SST-VIP comparison; p<0.001 for SST-EC comparison). VIP-expressing interneurons and excitatory cells did not show a significant difference (Mann-Whitney test, p=0.64). **(F)** 16 of 26 PV interneurons, 10 of 18 SST interneurons and only 1 of 12 VIP interneurons received greater synaptic inputs from S2 than ECs as they were above the 95 % CI (dashed blue) of the estimated eEPSC (solid blue) of ECs (See Methods; Fig. 1). **(G)** Comparison of synaptic weights onto each interneuron type for synaptic inputs from S2. Synaptic input bias values were calculated using the formula: (eEPSC in target inhibitory neuron − eEPSC in the excitatory cell) / (eEPSC in target inhibitory neuron + eEPSC in the EC). Input bias ranges between 1 and −1, with > 0 indicating the synaptic input is biased toward interneurons compared with EC in nwS1. Mean ± SEM values were; PV: 0.839 ± 0.060 (10 cells); SST: 0.535 ± 0.108 (10 cells); VIP: 0.118 ± 0.240 (9 cells). Input bias for the S2→nwS1 pathway was significantly greater than zero in PV and SST neurons, whereas VIP neurons displayed bias values close to zero (PV: p = 0.002, SST: p = 0.004, VIP: p = 0.734; Wilcoxon signed-rank test of the null hypothesis that mean input bias = 0). **p < 0.01.

**Figure 7.**
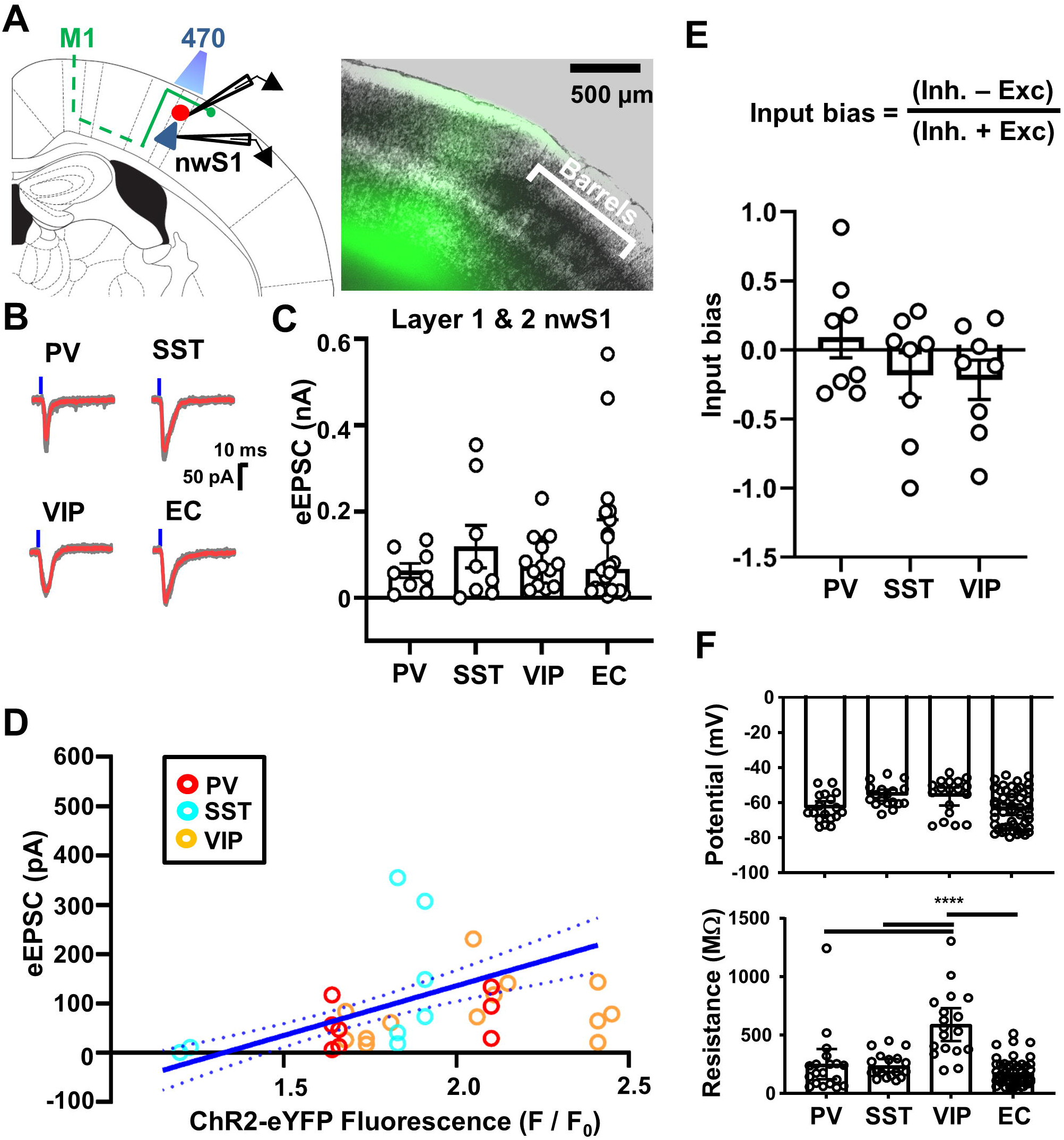
M1→nwS1 input broadly engages different subtypes. **(A)** Left: Schematic of slice recording configuration. An interneuron and a nearby putative excitatory neuron were recorded sequentially in pairs. Right: Example of a slice fixed in 10% formalin after recording showing expression of ChR2 in M1 after stereotaxic injection of AAV5- CamKIIa-hChR2(H134R)-EYFP to M1 (1.0 mm anterior, 1.0 mm lateral from bregma). **(B)** Representative response traces from sequential paired recordings on slices expressing tdTomato in PV-, SST-, or VIP-positive interneurons and excitatory cells (EC). Averages from five consecutive sweeps of optogenetically evoked EPSCs (eEPSCs, gray) are shown in red. Blue vertical lines indicate photostimulation (470 nm LED, 5 ms) delivered through the objective. Responses were acquired in the presence of tetrodotoxin and 4-aminopyridine. **(C)** eEPSC amplitudes of individual neurons in response to photostimulation of the M1→nwS1 projection. Each dot represents an average of five sweeps. Bar indicates mean; error bars indicate SEM. Mean ± SEM of peak eEPSC amplitude for PV-, SST-, VIP-expressing interneurons and ECs were 62.9 ± 16.9 pA (8 cells), 119 ± 49.3 pA (8 cells), 83.2 ± 17.1 pA (13 cells) and 119 ± 29.1 pA (24 cells) respectively. The M1 projection did not show preferential innervation of any particular subtypes (Mann-Whitney test, p > 0.1 for each comparison). **(D)** 1 of 8 PV neurons, 2 of 8 SST neurons and 1 of 13 VIP neurons were above the 95 % CI (dashed blue) of the estimated eEPSC (solid blue) of ECs (See Methods). **(E)** Comparison of synaptic weights onto each interneuron type for synaptic inputs from M1. Mean ± SEM values were; PV: 0.094 ± 0.152 (8 cells); SST: -0.183 ± 0.0.163 (8 cells); VIP: - 0.216 ± 0.142 (8 cells). Wilcoxon signed-rank test of the null hypothesis that mean input bias = 0. **(F)** Resting membrane potential (top) and input resistance (bottom) of all neurons included in the present study. ****p < 0.001, ANOVA with Tukey’s test.

A caveat with this interpretation is that variable hChR2 expression may contribute to the observed differences in eEPSC amplitudes. To control for the variability in hChR2 expression, we determined the line of best fit for the relationship between the eEPSC amplitude of excitatory cells and the expression of hChR2-EYFP across all brain slices used for recording (see Methods; Fig. 1). We then asked if eEPSC amplitude values of each subtype are significantly above or below 95% confidence interval (CI) of this linear model. About 62% of PV interneurons (16 of 26) and 55% of SST (10 of 18) interneurons were above the 95 % CI in contrast to only 1 of 12 VIP interneurons (Fig. 6*F*). To further control for variability in hChR2 expression, we also analyzed data from a subset of mice in which recordings were made sequentially from pairs of inhibitory and excitatory cells located within 50 µm of each other in the same slice (Fig. 6*G*). Initially we tried normalizing eEPSC amplitudes of the inhibitory neuron to those of the excitatory cell in each pair, which is standard practice in the field (Lee et al., 2013). However, some of the recorded pairs contained excitatory cells that were not responsive at all, which made normalization impossible. Instead, we calculated ‘input bias’, which quantifies the relative strength of synaptic input onto a particular inhibitory subtype over excitatory cells (See Methods; Fig. 6*G*). This metric is analogous to the ‘selectivity index’ widely used in sensory neuroscience to quantify the extent to which a neuron’s response is biased towards a certain sensory feature. The mean input bias values were; PV: 0.839 ± 0.060; SST: 0.535 ± 0.108; VIP: 0.118 ± 0.240.

Input bias for the S2→nwS1 pathway was significantly greater than zero in PV and SST neurons, whereas VIP neurons displayed bias values close to zero (PV: p = 0.002, SST: p = 0.004, VIP: p = 0.734; Wilcoxon signed-rank test). These results show that the S2 projection preferentially targets inhibitory PV and SST interneurons over excitatory cells and VIP interneurons in nwS1.

To test whether the effect on inhibitory interneurons is specific to the S2→nwS1 pathway or a general feature found of any cortico-cortical input in nwS1, we examined synaptic connections in the M1→nwS1 pathway using the same approach (Fig. 7). eEPSC recordings were targeted to layer 1 and layer 2 to match the spatial distribution of relevant axonal fluorescence (Fig 4). We photostimulated ChR2-expressing M1 axon terminals in nwS1 while recording eEPSCs in the three major interneuron subtypes as well as neighboring ECs (Fig. 7*A, B*). Data was expressed as the mean ± SEM of peak eEPSC amplitude. Values for PV-, SST-, VIP-expressing interneurons and ECs were 62.9 ± 16.9 pA (8 cells), 119 ± 49.3 pA (8 cells), 83.2 ± 17.1 pA (13 cells) and 119 ± 29.1 pA (24 cells) respectively (Fig. 7*C*). In contrast to the S2→nwS1 pathway, the M1 projection did not preferentially innervate any particular subtypes (p > 0.1 for each comparison, Mann-Whitney test). We compared eEPSC amplitude values of different subtypes against the linear model that links eEPSC amplitude of ECs to hChR2 expression level. In contrast to S2→nwS1 pathway, only 1 of 8 PV neurons, 2 of 8 SST neurons and 1 of 13 VIP neurons were above the 95% CI range in M1→nwS1 pathway (Fig. 7*D*). Input bias values calculated from inhibitory-excitatory cells in the same slice were not significantly greater than zero (p > 0.1 for each comparison, Wilcoxon signed-rank test; Fig. 7*E*). Resting membrane potential and input resistance of all neurons in the present study are shown in Figure 7F. Consistent with previous studies (Lee et al., 2013), VIP interneurons showed significantly higher input resistance than other subtypes (p < 0.001, ANOVA with Tukey’s test). From these results, we conclude that the S2 projection preferentially targets PV and SST interneurons in nwS1, whereas M1 projection broadly innervates all tested subtypes including VIP interneurons and excitatory cells.

### Loss of PV interneurons in nwS1 impairs motor coordination

PV interneurons in nwS1 receive a prominent synaptic input from S2. If PV interneurons are important for S1–S2 interaction during locomotion, inactivating PV neurons is expected to impair locomotion. Cre-dependent diphtheria toxin receptor (DTR) was expressed unilaterally in nwS1 by injection of adeno-associated virus (AAV1-flex-DTR-GFP) into PV-Cre; tdTomato^fl/+^ mice (Azim et al., 2014). We then injected these mice with intraperitoneal (i.p.) diphtheria toxin (DTX) and tested their performance on the rotarod (Fig. 8*A*). The ablation of tdTomato-labeled PV neurons was localized to nwS1 of mice that received DTX (Fig. 8*B*). A separate group of mice expressing DTR received saline injections. DTX-injected mice showed a shorter ‘latency to fall’ on average than saline-injected mice (Fig. 8*C*). Rotarod performance was significantly impaired in DTX-treated mice (saline: 212 ± 6.86 s, 7 mice; CNO: 159 ± 2.64 s, 8 mice; t(13) = 3.65, p = 0.0029, *t*-test, g = 1.89).

**Figure 8.**
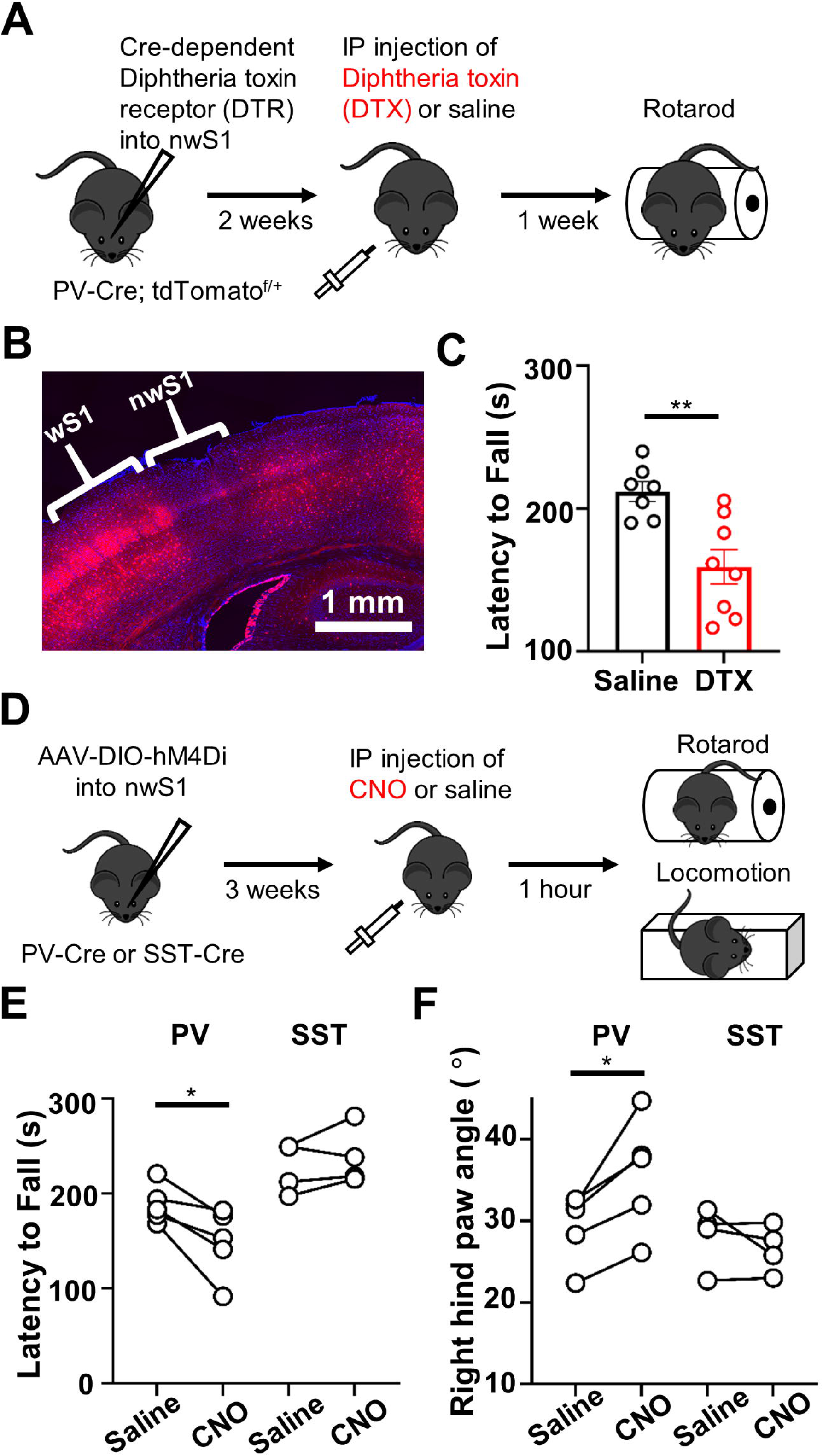
Inactivation of PV neurons in nwS1 causes deficits in locomotion. **(A)** Experimental paradigm. Cre-dependent diphtheria toxin receptor (DTR) was expressed in nwS1 PV neurons via stereotaxic virus injection into mice expressing Cre in PV neurons (coordinates from bregma: posterior, 0.7 mm; lateral, 2.5 mm). Two weeks after injection, mice received intraperitoneal diphtheria toxin (DTX; 30 ng/g) or an equivalent volume of saline once daily for 2 days. One week later, the mice were subjected to the accelerated rotarod assay. **(B)** Ablation of tdTomato-labeled PV neurons (red) in nwS1 after unilateral injection of AAV1- Flex DTR-IRES-GFP and i.p. injection of diphtheria toxin. PV neurons in nwS1 were selectively ablated, whereas those in wS1 were spared. **(C)** Average rotarod performance of mice in the DTX and saline groups. Saline: 212 ± 6.86 s, 7 mice; DTX: 159 ± 12.1 s, 8 mice; ** p=0.0029, *t*-test. **(D)** Experimental paradigm. AAV9-hSyn-DIO-hM4D(Gi)-mCherry was injected into nwS1 of PV-Cre or SST-IRES-Cre mice. Three weeks later, mice received an intraperitoneal injection of CNO (CNO-2HCl, 4.13 mg/kg) or the equivalent volume of saline 1 h prior to testing on the rotarod and the locomotion assay. **(E)** Average rotarod performance of PV- and SST-Cre mice with saline and CNO injections. PV inactivation; saline: 189 ± 9.05 s; CNO: 149 ± 16.1 s, 5 mice; p=0.022, paired *t*-test. SST inactivation; saline: 227 ± 13.4 s; CNO: 238 ± 15.2 s, 4 mice; p=0.306, paired *t*-test. **(F)** Mean right hind paw angles in mice before and after the PV or SST inactivation. Each data point represents a mean of the hind paw angles during multiple strides observed in a single mouse. PV inactivation; saline: 29.4 ± 1.90; CNO: 35.7 ± 3.14; 5 mice; p=0.019, paired *t*-test. SST inactivation; saline: 28.2 ± 1.89; CNO: 26.6 ± 1.45; 4 mice; p=0.332, paired *t*-test.

While this result suggests that PV interneurons in nwS1 regulate locomotion by interfacing with S2 inputs, inactivation of any interneuron subtypes in nwS1 might impair locomotion. To test whether S2 → PV is specifically required for the control of paw movement during locomotion, we assessed the impact of silencing the SST population, which also receives a major synaptic input from S2. AAV encoding Cre-dependent hM4Di-mCherry was injected into nwS1 of PV-Cre or SST-Cre mouse. Three weeks after hM4Di expression, mice received an i.p. injection of CNO (4.13 mg/kg) or saline and were tested on the rotarod and the runway (Fig. 8*D*). The CNO-injected PV-Cre mice showed significant impairments in rotarod performance with a shorter average ‘latency to fall’ time than the saline-injected group (saline: 189 ± 9.05 s; CNO: 149 ± 16.1 s, 5 mice; t(4) = 3.63, p = 0.022, g = 1.36, paired *t*-test; Fig. 8*E*). On the other hand, silencing the SST population did not affect rotarod performance; differences between the CNO and saline-treated groups were not observed in these mice (saline: 227 ± 13.4 s; CNO: 238 ± 15.2 s, 4 mice; t(3) = 1.23, p = 0.306, g = 0.384, paired *t*-test; Fig. 8*E*). The runway locomotion assay revealed that PV inactivation increased the right hind paw angle at the onset of swing phase compared with saline injection in the same mice (saline: 29.4 ± 1.90; CNO: 35.7 ± 3.14; 5 mice; t(4) = 3.78, p = 0.019, g = 1.50, paired *t*-test; Fig. 8*F*). In contrast, SST inactivation did not alter the hind paw angle (saline: 28.2 ± 1.89; CNO: 26.6 ± 1.45; 4 mice; t(3) = 1.15, p = 0.332, paired *t*-test, g = 0.475; Fig. 8*F*). Based on these results, we conclude that the S2 → PV pathway in nwS1 is critically involved in maintaining hind paw orientation during locomotion. Note that neither DTR nor DREADD-mediated manipulation used in the present study was specifically targeted to those PV neurons that receive S2 input. Future studies will test whether PV neurons receiving S2 input can be selectively ablated by combining the Flp- and Cre-dependent DTR and anterograde trans-synaptic virus in PV-Cre background.

In summary, our results demonstrate that S2 input and local PV interneurons in nwS1 are critically involved in the regulation of efficient paw orientation during locomotion. We also report that different long-range cortico-cortical inputs engage distinct interneuron subtypes as a postsynaptic target. Together, these findings expand our understanding of cortical feedback processing by bridging the local circuit organization and function of different cortico-cortical connections in the same target area.

## Discussion

Our data provide insight into how specific cortico-cortical connections in the primary somatosensory cortex regulate the complex process of locomotion. The importance of somatosensory input in locomotion has previously been demonstrated. For example, genetic ablation of proprioceptive afferents in the periphery is known to disrupt locomotion (Chen et al., 2007, Akay et al., 2014). ‘Sensory ataxia’ is indeed a hallmark of peripheral neuropathies, such as Charcot-Marie-Tooth disorder. On the other hand, how central somatosensory processing contributes to rhythmic movement like locomotion has remained elusive. Here we demonstrate that the S1 cortex is required for regulating hind paw movements during locomotion. The cortico-cortical projection originating from S2 participates in the control of locomotion, whereas the M1 projection does not. The differential involvement of S2 and M1 inputs implies that they provide different information and/or innervate different targets in the S1 circuit. The latter is supported by our results showing that the two cortical pathways also differ in terms of their postsynaptic targets in S1. The S2 projection preferentially targets inhibitory interneurons over excitatory neurons, whereas the M1 projection shows broader target engagement. Collectively, our results show that S2→S1 pathway is required for maintaining efficient movements of the paw by regulating its orientation during locomotion. This regulation potentially involves PV and/or SST interneuron populations in the local circuit.

### The role of S1 in the control of paw movement during locomotion

Although locomotion is typically considered a highly autonomous motor activity regulated by brain stem-spinal cord circuits, the role of the cerebral cortex in the planning and execution of locomotion has been demonstrated in studies across several different species (for a recent review, see Drew and Marigold, 2015). Modulation of S1 neuronal activity correlates with locomotor output in primates (Fitzsimmons et al., 2009), cats (Favorov et al., 2015) and rodents (Chapin and Woodward, 1982a). Kinematic parameters of walking can be decoded from the activity of S1 neurons (Fitzsimmons et al., 2009). More recently, a direct role of S1 in controlling locomotion has also been demonstrated using causal manipulation in mice. Chemogenetic activation (or inhibition) of S1 activity accelerated (or impaired) locomotion (Karadimas et al., 2020). However, specific aspects of locomotion regulated by S1 remained uncharacterized. During locomotion, S1 can directly generate a motor response, encode somatosensory information or serve both of these functions.

Our results demonstrate that the unilateral inactivation of S1 increased the degree of paw slip on the rotarod and altered the hind paw angle during walking (Fig. 2). We argue that these impairments are caused by disruptions in somatosensory feedback during locomotion rather than the inability to generate motor output. First, the inactivation of S1 did not influence mouse’s ability to initiate and complete the corrective step during the rotarod performance. Second, the S1 manipulation did not affect the speed of locomotion, which typically decreases when M1 is inactivated (Warren et al., 2021). We propose that movement-related somatosensory feedback encoded in S1 is important for maintaining the orientation of paw movements during locomotion. Future studies will uncover the precise nature of information encoded in S1 (cutaneous or proprioceptive) that is utilized by downstream locomotor centers.

### The role of long-range cortico-cortical projections in locomotion

We speculate that the S2→S1 pathway may contribute to locomotion in different ways. First, S2 input may enhance the fidelity of somatosensory feedback during movement by sharpening feature coding in the S1. In the mouse whisker S1 (wS1), silencing S2 was shown to degrade orientation selectivity, indicating that the S2 activity enhances orientation tuning in wS1 (Minamisawa et al., 2018). Likewise, the orientation selectivity of neurons in the mouse primary visual cortex (V1) decreased when the secondary visual cortex (V2) was silenced (Pafundo et al., 2016). The V2→V1 pathway also contributes to surrounding suppression (Nurminen et al., 2018, Vangeneugden et al., 2019). Neural components of cortico-cortical connections that participate in the modulation of feature selectivity and surround suppression remain to be determined. Nevertheless, it is becoming clear that higher→lower sensory cortical connections like S2→S1 (Fig. 6) and V2→V1 pathways innervate PV and SST inhibitory interneurons (Gonchar and Burkhalter, 2003, D’Souza et al., 2016, Wall et al., 2016). GABAergic inhibition is crucial for shaping feature tuning and surround suppression in sensory cortex (Nelson et al., 1994, Katzner et al., 2011, Adesnik, 2017). Therefore, a potential role of the S2→S1 pathway is to increase the fidelity of somatosensory feedback during movement by enhancing feature selectivity and suppressing responses to irrelevant stimuli. This possibility can be tested by characterizing somatosensory tuning of nwS1 neurons while manipulating the S2→S1 pathway. Second, S2 may transmit movement-related signals such as corollary discharge, predicted sensory consequences and the context of movement to the S1 (Kwon et al., 2016, Condylis et al., 2020). S2 is reciprocally connected with M1 (Suter and Shepherd, 2015), suggesting that the corollary discharge signal generated in M1 may reach S1 through S2. These movement-related signals may be important for filtering out sensory responses generated during self-movement. This has been shown in the auditory system (Schneider et al., 2014) and for computing sensory prediction errors during motor adaptation (Mathis et al., 2017).

Silencing M1→nwS1 did not impact rotarod performance or over-ground locomotion. While the reason for the lack of the effect is unclear, there are several possible explanations. The stereotaxic coordinates used for injecting Cre-dependent hM4Di into M1 were based on the location of nwS1-projecting M1 neurons in the retrograde tracing experiment (Fig. 3*B*) and may not correspond to a hind paw M1 area. It is unknown to what extent the areas defined using anatomical tracing and functional mapping are aligned with each other. Retrograde labeling of M1 neurons projecting to nwS1 and intracortical microstimulation should be combined in the same animal to address this issue. It is also possible that while M1 itself is crucial for baseline locomotion (Warren et al., 2021) or that the M1→nwS1 pathway is involved in a different function such as motor learning (Kawai et al., 2015). Lastly, the simple motor assays used in the present study may be insufficient to capture complex changes in paw kinematics caused by silencing of M1→nwS1 connections.

### Synaptic organization of long-range cortico-cortical inputs in S1

The S2→nwS1 connection provides direct synaptic inputs to PV and SST inhibitory interneurons. M1→nwS1 inputs are distributed among all major interneuron subtypes. Recent studies have suggested that long-range cortical projections recruit VIP interneurons that inhibit other interneurons, leading to the disinhibition of pyramidal neurons (Lee et al., 2013, Pfeffer et al., 2013, Pi et al., 2013, Fu et al., 2014, Karnani et al., 2016). Our results show that different projections engage different neuronal subtypes as a postsynaptic target. VIP neurons are the primary target of M1→wS1 input and mediate disinhibition in the local circuit (Lee et al., 2013). Other studies support a broader engagement of excitatory and inhibitory subtypes in the M1→wS1 pathway (Kinnischtzke et al., 2014, Kinnischtzke et al., 2016). Our observation that the M1→nwS1 pathway innervates all three interneuron subtypes with little bias provides additional support for these latter studies. Note that wS1 and nwS1 may differ in terms of the composition and spatial distribution of interneuron subtypes. The VIP pool may consist of different subsets, including neurons that are exclusively involved in M1→S1 pathway.

While our results demonstrate that the S2 and M1 pathways are functionally distinct, how these pathways influence S1 activity was not examined in the present study. What would the net effect of activating or inactivating each of these pathways be? Given the prevalence of recurrent connections in the neocortex, the outcome of activating different interneuron subtypes is not intuitive, and is perhaps even paradoxical (Fu et al., 2014, Rubin et al., 2015, Litwin-Kumar et al., 2016). The net effect of activating the S2→nwS1 or M1→nwS1 pathway could be facilitation or suppression of output spiking in nwS1 pyramidal neurons; a positive vs. negative effect may depend on a number of factors such as baseline activity, operating regime of the network, number of activated neurons and the strength of sensory stimuli (Rubin et al., 2015, Garcia Del Molino et al., 2017, Sadeh et al., 2017). This is an important question for future studies. Nevertheless, our findings suggest that differences in the postsynaptic target engaged by different long-range cortico-cortical connections should be taken into consideration when constructing theoretical models of cortical networks.

In summary, we demonstrate that non-whisker S1 is critical for maintaining the orientation of paw movement during locomotion and that the S2 input is important for this process. While S2→S1 pathway engages both PV and SST interneuron subtypes as a postsynaptic target, PV cells are directly involved in the control of paw movement. Altogether, we establish a causal role for S1 in regulating hind paw movements during locomotion and elucidate components of the cortico-cortical circuit involved in this process.

## Acknowledgments

We thank members of the Kwon laboratory, Richard Hume and Daniel O’Connor for critical reading of the manuscript, and Bo Duan’s lab for technical assistance with rotarod. This work was supported by a Bridge to Independence grant from the Simons Foundation Autism Research Initiative (S.E.K).

